# LNetReduce: tool for reducing linear dynamic networks with separated time scales

**DOI:** 10.1101/2021.05.11.443578

**Authors:** Marion Buffard, Aurélien Desoeuvres, Aurélien Naldi, Clément Requilé, Andrei Zinovyev, Ovidiu Radulescu

## Abstract

We introduce LNetReduce, a tool that simplifies linear dynamic networks. Dynamic networks are represented as digraphs labeled by integer timescale orders. Such models describe deterministic or stochastic monomolecular chemical reaction networks, but also random walks on weighted protein-protein interaction networks, spreading of infectious diseases and opinion in social networks, communication in computer networks. The reduced network is obtained by graph and label rewriting rules and reproduces the full network dynamics with good approximation at all time scales. The tool is implemented in Python with a graphical user interface. We discuss applications of LNetReduce to network design and to the study of the fundamental relation between time scales and topology in complex dynamic networks.

**Availability:** the code and application examples are available at https://github.com/oradules/LNetReduce.

## 1 Introduction

In bioinformatics and systems biology, molecular networks are used as mechanistic models of cell physiology and disease with numerous applications in biology and medicine. Networks are also used by the complex systems community to study social interactions, epidemics, or computer communication. In various fields, large scale networks are available as digraphs, in which vertices and edges represent individuals (for instance molecules) and interactions, respectively. Network topology is supposed essential for their properties, therefore a large number of tools are dedicated to the analysis of network topology [4]. However, network dynamics is also very important. The simplest model of dynamic network is obtained by associating to each edge, a number representing the strength of the interaction or its time scale. For molecular networks, this type of information can result from quantitative network analysis approaches such as modular response analysis, flux balance analysis, or from direct probing of the interaction by biochemical or biophysical methods. When interaction time scales are not accurate, one can represent their values by integer orders of magnitude instead of real numbers. In many cases, it is important to know that one interaction is much faster than another without having to know by precisely how much. Integer labelled digraphs are thus well suited to study network properties that depend on timescale orders. In this paper we introduce a tool to simplify such networks to an extent that qualitative analysis of their dynamics becomes easy.

## 2 Model

Dynamic networks are represented as integer edge-labeled digraphs 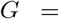 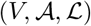, with *V* the set of vertices, 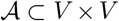 the set of edges, 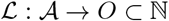 the label function. The labels can be obtained from timescales as follows. Fixing a scale basis 0 < *ϵ* < 1, to each edge with kinetic constant *k* and time scale *τ* = 1/*k* and we associate an integer kinetic order *g* = *round*(log(*k*)/ log(*ϵ*)). *g* is the order of magnitude of *k*, as shown by *ϵ*^g−0.5^ ≤ *k* < *ϵ*^g+0.5^. The time units are chosen such that the fastest reaction has *k* = 1, *g* = 0, which results in positive integer orders. When *ϵ* = 1/10 one recovers the familiar decimal orders of magnitude. Using this power parametrisation we can cope with widely distributed rates. Such networks can be endowed with deterministic or stochastic dynamics.

The deterministic dynamics is defined by a set of ODEs:

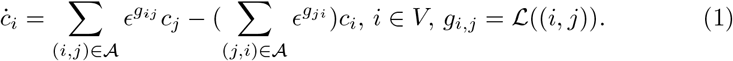

The stochastic dynamics is a random walk on the network, where the probability to jump from *i* to *j* is proportional to **ϵ**^*g*_*ji*_^. For continuous time random walks, (1) is the backward Kolmogorov equation (master equation) and *c*_*i*_ is the probability to be in *i*.

## 3 Reduction algorithm

We are interested in the reduced model valid in the limit *ϵ* → 0. This model can be obtained algorithmically using the following rules [3, 6, 5, 7]:

1. **Pruning**. For any node with several successors keep the edge with minimum order *g*_*i*_ = min{*g*_*ij*_, *j ϵ Succ*(*i*) and delete all the other edges. We ask for the **condition 1**: *at a bifurcation the minimum order is attained only once.* The result of this step is the deterministic auxiliary network *Aux*(*G*). If the auxiliary network is acyclic, the algorithm stops after this step.
2. **Pooling**. If *Aux*(*G*) contains cycles, find a maximal set of disjoint irreducible cycles. Replace these cycles by “glued nodes”. As pooling will eventually apply several times, it generates hierarchical glued nodes. Each glued node retains the memory of the cycles it contains (nodes and edges) as follows:

– The glued node inherits all the edges of *Aux*(*G*) entering the cycle and also all the edges of *G* exiting the cycle.
– The labels of edges from *Aux*(*G*) are maintained, while the labels of edges of *G* and not in *Aux*(*G*) are recomputed according to the rule:

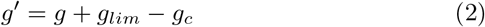

where *g′, g* is the order after and before gluing, respectively, *g*_*lim*_ is the largest order edge in the cycle (limiting step) and *g*_*c*_ corresponds to the cycle edge sharing the tail with the exit edge. Here we ask for the **condition 2:** *the limiting step is unique in all cycles*.

If the application of pooling results in a non-deterministic graph, apply pruning again. Iterate until there are no more cycles.

3. **Restore** glued vertices through the following steps:

– Restore all vertices of the glued cycles.
– Restore all cycle edges except the limiting step.
– An edge exiting the glued cycle and arriving in an unglued node, is replaced by an edge with the same head and label, but originating from the tail of the limiting step.
– An edge exiting the glued cycle and arriving in another glued cycle is replaced as above using its original head within the glued cycle.

The result of restore is path independent: one can start with the most compact or less compact cycles in the hierarchy.

The top level version of the algorithm is given by Algorithm 1. If conditions 1 and 2 are everywhere satisfied, then the reduced graph is acyclic and deterministic. Exceptions lead to stopping the reduction before eliminating all cycles and multiple branching. An example of application of the algorithm to “flower” motifs, consisting of a central hub node and satellite nodes is shown in Figure 1.

**Fig. 1.**
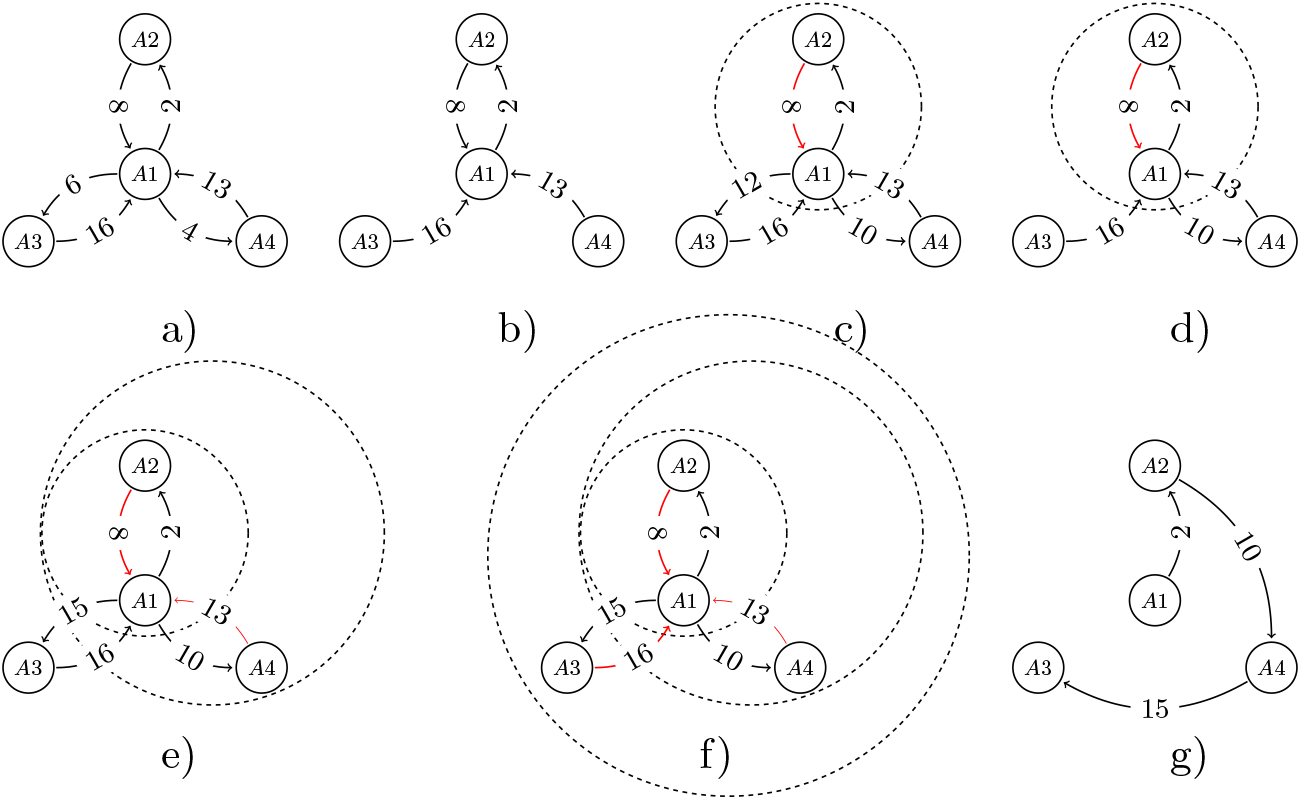
The successive steps of the reduction algorithm. a) is the initial model; b) is the auxiliary network resulting from pruning; c) is the result of gluing the cycle {*A*_1_, *A*_2_} and rewriting the exit edge labels (the labels 6, 4 become 6 + 8 − 2 = 12, and 4 + 8 − 2 = 10, respectively); d) is the auxiliary network after one more iteration; e) results from gluing the cycle {{*A*_1_, *A*_2_}, *A*_4_} and rewriting the exit edge label (12 becomes 12 + 13 − 10 = 15); f) results from gluing the cycle {{{*A*_1_, *A*_2_}, *A*_4_}, *A*_3_}; g) restoring the single species without their limiting steps starting with the innermost cycle. Limiting steps of different cycles are represented in red.

### Algorithm 1 Reduce

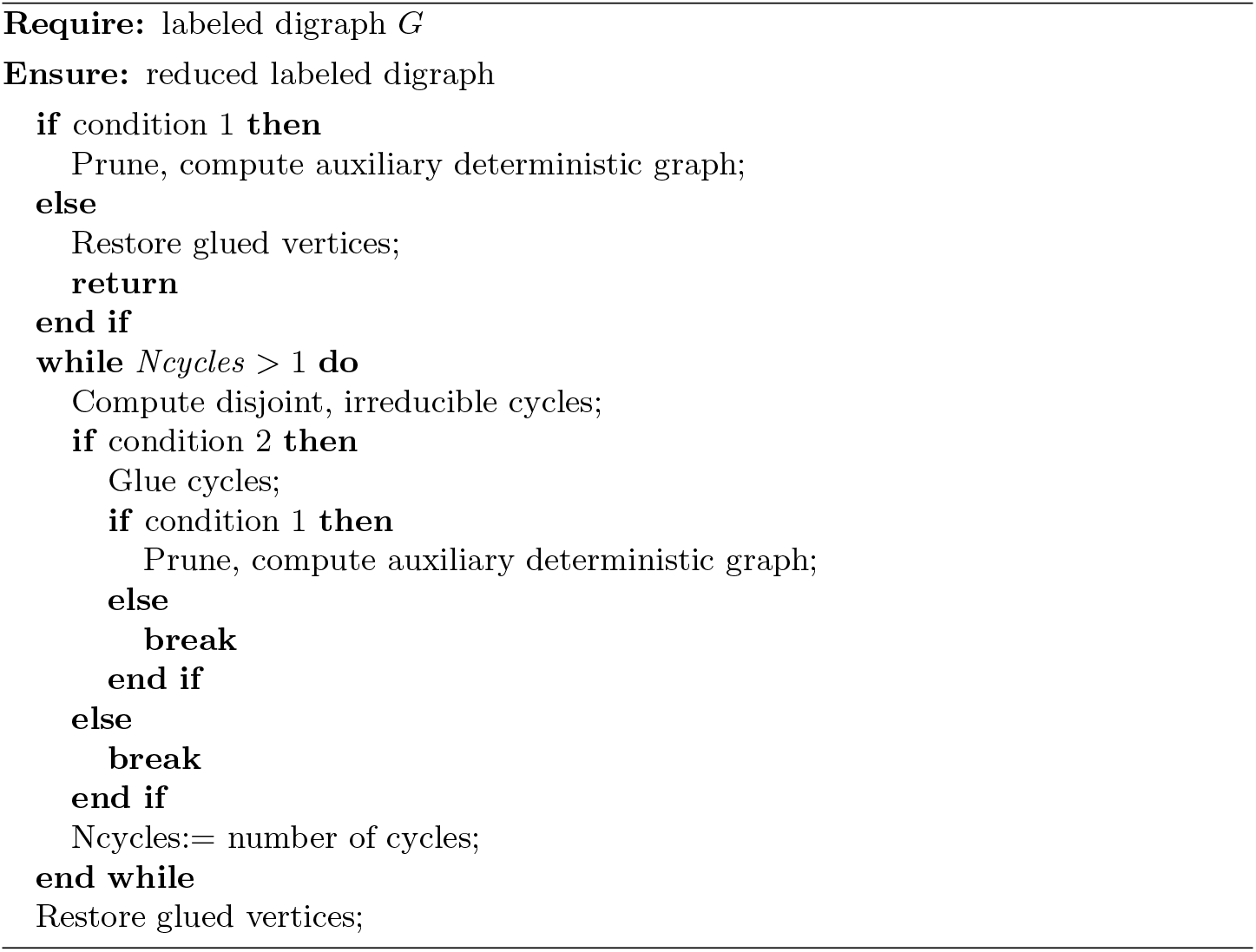

## 4 Applications

### 4.1 Connection between topology and dynamics

For a given topology one can have several reduced models, depending on the kinetic orders. Each reduction corresponds to a particular qualitative dynamics.

In order to illustrate the connection between network topology and its dynamics in the limit of well-separated rate constants, we made a number of experiments on fragments of real-life transcription networks extracted from Dorothea database (https://saezlab.github.io/dorothea/), for the edges of which we assigned sequential and distinct kinetic orders. For example, we extracted all network neighbours of MYCN transcription factor (see Figure 2,A, and randomly assigned kinetic orders from 0 to 9, to network reactions. Application of LNetReduce to 10000 random kinetic order assignments led to 51 topologically distinct (topologically isomorphic with node identity kept) model reductions, where some reductions were much more frequent than others. Of note, in approximately 7% cases, LNetReduce met a conflict in the reduction algorithm. In these cases, two kinetic rates of the same order in the outgoing reaction fork happened at a stage of reduction process. Interestingly, rare reductions (Figure 2C,D) were typically characterized by the presence of reactions with effectively opposite direction compared to the initial network, and inefficient, leaky dynamics of MYCN, with small probability of visiting it by a random walk at longer timescales (as opposed to the activating and efficient dynamics of the most frequent reductions, Figure 2A,B). Other computation experiments with fragments of the transcription network are described at the LNetReduce web-site as Python notebooks.

**Fig. 2.**
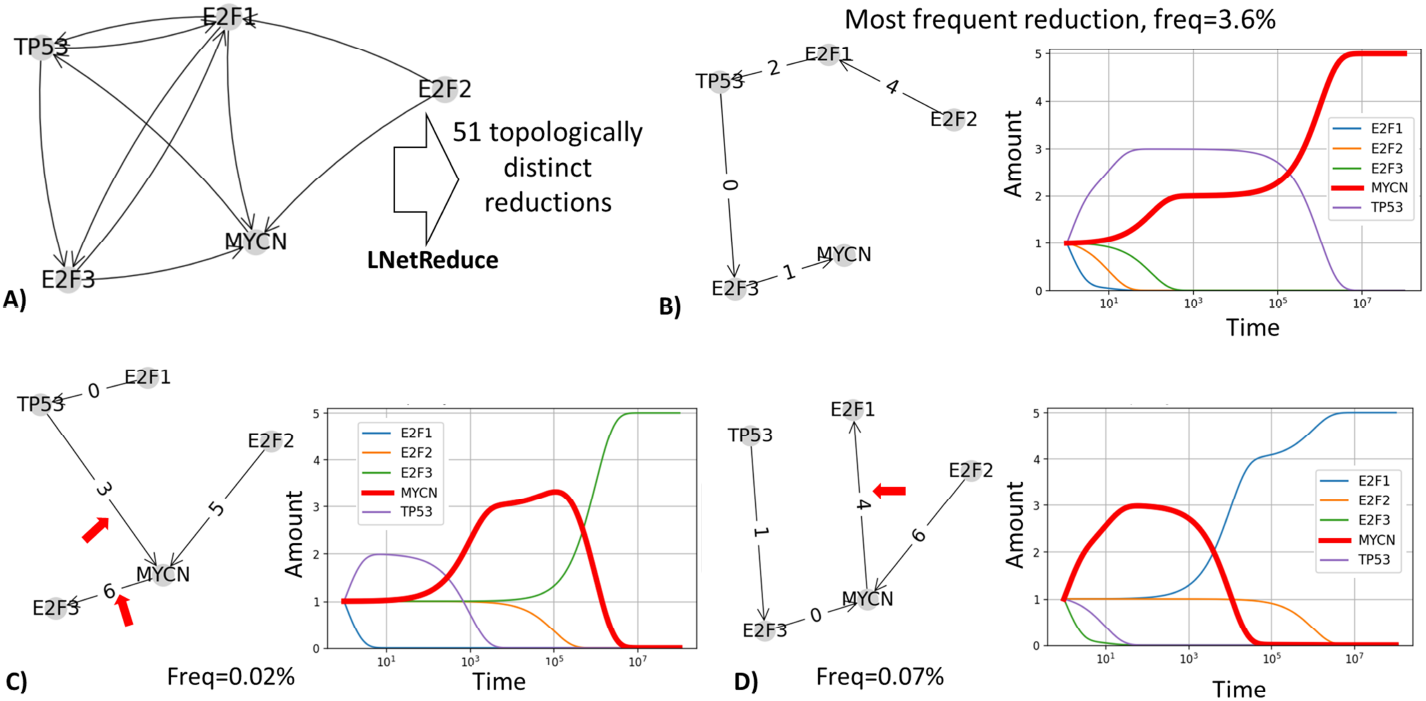
Example of LNetReduce application for studying the connection between the network topology and dynamics, using a small fragment of experimentally obtained transcription regulation network. A) Network fragment. B) Most frequent topologically isomorphic reduction and its dynamics. C,D) Examples of rarely obtained reductions and their dynamics. Red arrows show edges whose direction is reverted with respect to A).

### 4.2 Design of slow transients

The reduced graph is a forest of inverted trees. For such a model, relaxation time scales defined as times after which something happens are simply the new labels [3]. However, the relation between initial step time scales *g*_*ij*_ and the final labels can be intricate. In particular, relation (2) shows that timescales tend to be larger than timescales of the initial steps and complex networks tend to have very slow transients. Slow transients are important in biology for a variety of processes from cellular memory to long period circadian oscillations. We can formulate the following design rule for long transients:

#### Rule 1

*At least at some iteration, the auxiliary network must contain cycles such that the cycle step at an exit point is not the limiting step, *g*_*lim*_ > *g*_*c*_. Then, according to (2) *g′ > g*, new steps are slower than older*.

Rule 1 is responsible for the slow transient in Figure 3. This rule also explains the counter-intuitive property that a non-uniform increase of all switch rates in a stochastic network can lead to slower transients [2]. Large networks can more easily generate slow transients by rule 1 or by simple addition, in the presence or absence of separation, respectively. This explains why proteins controlling circadian clock in *Neurospora* have about one hundred phosphorylation sites [1].

**Fig. 3.**
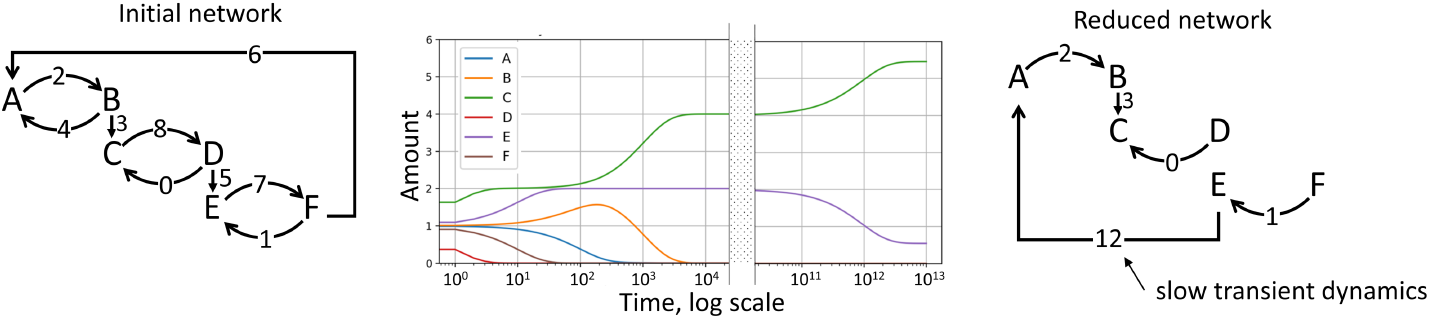
Application of LNetReduce to identify slow transient dynamics in networks. The cascade shown on the left, after fast initial dynamics achieves a very slowly relaxing state (middle panel). The timescales of kinetic rates are shown as numbers on reaction arcs (the smaller the faster). On the right the result of application of LNetReduce is shown, explicitly revealing the existence of a very long timescale in the network, four orders of magnitude larger than any reaction in the initial network (12 vs 8). Using random permutations of kinetic orders, we estimate the frequency of emergence of slow transients with this network topology to 4%.

### 5 Conclusion

We provide a tool allowing to study dynamics of networks with separate timescales. The separation criteria are formulated as the the Rules 1,2 in the paper. The tool computes a reduced model that is a forest of inverted trees. The reduced model provides immediately the relaxation timescales of the network and its qualitative dynamics. For random walk applications, our tool represents an alternative to uniform switching rates. For network design it allows to study the interplay between time scales and topology for predicting network dynamics and eventually controllability. In future work we will consider situations when some of the separability conditions can be released.

## 6 Acknowledgements

This work was supported by Agence Nationale de la Recherche, projects ANR-17-CE40-0036 SYMBIONT and ANR-19-P3IA-0001 (PRAIRIE 3IA Institute), and by the Ministry of Science and Higher Education of the Russian Federation (project No. 14.Y26.31.0022).

## References

1. Baker, C.L., Kettenbach, A.N., Loros, J.J., Gerber, S.A., Dunlap, J.C.: Quantitative proteomics reveals a dynamic interactome and phase-specific phosphorylation in the neurospora circadian clock. Molecular cell 34(3), 354–363 (2009)

2. Bokes, P., Klein, J., Petrov, T.: Accelerating reactions at the dna can slow down transient gene expression. In: International Conference on Computational Methods in Systems Biology. pp. 44–60. Springer (2020)

3. Gorban, A.N., Radulescu, O.: Dynamic and static limitation in multiscale reaction networks, revisited. Advances in Chemical Engineering, Volume 34 (2008), 103–173 (Mar 2007). https://doi.org/10.1016/S0065-2377(08)00003-3

4. Hagberg, A., Swart, P., S Chult, D.: Exploring network structure, dynamics, and function using networkx. Tech. rep., Los Alamos National Lab.(LANL), Los Alamos, NM (United States) (2008)

5. Radulescu, O., Gorban, A.N., Zinovyev, A., Noel, V.: Reduction of dynamical biochemical reactions networks in computational biology. Frontiers in genetics 3, 131 (2012)

6. Radulescu, O., Gorban, A.N., Zinovyev, A.Y., Lilienbaum, A.: Robust simplifications of multiscale biochemical networks. BMC Syst. Biol. 2, 86 (2008). https://doi.org/10.1186/1752-0509-2-86

7. Radulescu, O., Samal, S.S., Naldi, A., Grigoriev, D., Weber, A.: Symbolic dynamics of biochemical pathways as finite states machines. In: Roux, O.F., Bourdon, J. (eds.) Computational Methods in Systems Biology - 13th International Conference, CMSB 2015, Nantes, France, September 16-18, 2015, Proceedings. Lecture Notes in Computer Science, vol. 9308, pp. 104–120. Springer (2015). https://doi.org/10.1007/978-3-319-23401-410, https://doi.org/10.1007/978-3-319-23401-4_10

